# *Wzc* and *WcaJ* mutations in hypervirulent *Klebsiella pneumoniae* lead to phage resistance at the cost of reduced virulence

**DOI:** 10.1101/2021.04.21.440873

**Authors:** Lingjie Song, Xianggui Yang, Jinwei Huang, Xiaokui Zhu, Guohui Han, Yan Wan, Ying Xu, Guangxin Luan, Xu Jia

## Abstract

Hypervirulent *Klebsiella pneumoniae* (hvKp) is one of the major community-acquired pathogens, which can cause invasive infections such as liver abscess. In recent years, bacteriophages have been used in the treatment of *Klebsiella pneumoniae*, but the characteristics of the phage resistant bacteria produced in the process of phage therapy need to be evaluated. In this study, two podoviridae phages, hvKpP1 and hvKpP2, were isolated and characterized. In *vitro* and in *vivo* experiments demonstrated that the virulence of the resistant bacteria was significantly reduced compared with that of the wild type. Comparative genomic analysis of monoclonal sequencing showed that nucleotide deletion mutations of *wzc* and *wcaJ* genes led to phage resistance, and the electron microscopy and mucoviscosity results showed that mutations led to the loss of the capsule, meanwhile, animal assay indicated that loss of capsule reduced the virulence of hvKp. The findings can contribute to a better understanding of that bacteriophage therapy can not only kill bacteria directly, but also reduce the virulence of bacteria by phage screening.

**Importance:** Bacteriophages are considered potential therapeutic alternative to antibiotics; however host-evolved phage resistance has accounted for the resurgences of pathogens, meaning further measures are need to improve the efficacy of phage therapy. This study showed two phages capable of infecting hypervirulent *K. pneumoniae* and identified phage-resistant mutants whose virulence was significantly reduced. Gene sequencing analysis revealed that mutations of *wzc* and *wcaJ* gene, related to capsule synthesis, recovered phage resistance but reduced the virulence of hypervirulent *K. pneumoniae*.

## Introduction

*Klebsiella pneumoniae*, a Gram-negative enteric bacterium, is one of the most important opportunistic nosocomial pathogens. Compared with the classic *K. pneumoniae* (cKp) strains that cause other types of opportunistic infections, *K. pneumoniae* strains that mainly cause pyogenic liver abscesses often have a substantially higher virulence and are therefore designated hypervirulent *K. pneumoniae* (hvKp). Hypervirulent *K. pneumoniae* has the ability to cause life-threatening, community-acquired infections, including liver abscesses complicated by endophthalmitis, meningitis, osteomyelitis, and necrotizing fasciitis, in young and healthy individuals and is therefore associated with high morbidity and mortality. In recent years, multidrug-resistant hypervirulent strains have emerged [1, 2], creating a new challenge in combating this already dangerous pathogen. Therefore, the therapeutic use of bacteriophage has garnered renewed attention [3] and is considered a proven strategy [4].

For phage therapy, one of the main research focuses is phage-resistant bacterial variants, that readily emerge in vitro [5], as the mechanisms of phage resistance, including preventing phage adsorption, preventing phage DNA entry, and cutting phage nucleic acids, are complex and varied [6]. The blocking of phage adsorption receptors is the most important evasive strategy for *K. pneumonia* [7, 8], of which capsules act as phage adsorption receptors in this bacterium, similar to *E. coli* and *A. baumannii* [9, 10].

Interestingly, capsular polysaccharide, a polysaccharide matrix that coats the cell, has been identified as a virulence factor [7]. In bacteria, the capsule confers resistance against the bactericidal activity of antimicrobial peptides, complement, and phagocytes [7, 8] and is synthesized by gene products from the capsular polysaccharide synthesis (cps) locus. In *K. pneumoniae*, the *cps* gene cluster has a total length of 21-30 kb, including 16-25 genes and harbors a number of genes involved in capsule production, including *wzi, wza, wzb, wzc, gnd, wcaJ, cpsB, cpsG*, and *galF*. Among them, the deletion or mutation of *wzi, wza, wzb* and *wzc* genes has a significant impact on virulence [11, 12], and enhances the phagocytosis of neutrophils and serum sensitivity of the strain [13]. In addition, *wbaP, wbaZ, wcaN, wcaJ, and wcaO* determine the diversity of capsule types, as reported in *E. coli* [14]. More than 70 capsular serotypes have been reported for *K. pneumonia*, of which K1, K2, K20, K54, and K57 are considered to be the main capsular types of hypervirulent strains [15]. To infect these hypervirulent strains, phages must be able to cross the capsule layer prior to docking and penetration. For this purpose, phages possess specific polysaccharide depolymerase enzymes, which degrade the polysaccharide capsule structure. However, these capsule depolymerases have specific effects and selectivity to certain serotypes [16]. In pursuit of identifying novel approaches for therapy, it is therefore important that relationships between phage resistance and virulence in the hvKps are explored.

In the present study, two bacteriophages against K57 capsular *K. pneumonia* were isolated from the sewage and in *vitro* and *vivo* treatment experiments were performed to evaluate the virulence of the mutant strains resistant to these two bacteriophages using the *G. mellonella* model. Sequence analysis revealed two genes, *wzc* and *wcaJ*, that had deletion mutations, and the role of these two genes in phage resistant mutant strains was studied.

## Materials and methods

### Bacteria strains and growth conditions

*Klebsiella pneumoniae* strains isolated from four hospitals in China were used in this study (Table 2). All strains were stored at -80 °C in 15% (v/v) glycerol, and all culturing was carried out in lysogeny broth (LB) at 37 °C with shaking at 200 rpm.

### Phage isolation

Phages were isolated from a local wastewater station in Chengdu, Sichuan. Briefly, untreated sewage was mixed with hvKpLS7 culture at a ratio of 1:1. The enrichment culture was then incubated overnight at 37 °C, centrifuged (5000RPM, 5min), and the supernatant filtered through a 0.22um filter to remove cells. This filtrate was mixed with hvKpLS7 in molten semisolid soft agar (0.7% agar) and poured over solidified 1.5% nutrient agar plates. All overlay agar plates were allowed to set, then incubated overnight at 37 °C. The resulting plaques were subjected to three rounds of plaque purification. The propagation of bacteriophages was determined according to the protocols of the Erna Li laboratory [17]. Purified phages were stored at 4 °C in SM buffer (100 mM NaCl, 8 mM MgSO_4·_7H_2_O, and 50 mM Tris-Cl (pH 7.5)), and the titer was determined using double-layer agar.

### Transmission electron microscopy of phage

Phage particles were spotted onto a carbon-coated copper grid and negatively stained with 2% (w/v) phosphotungstic acid (PTA). After drying, phages were observed on a Tecnai G^2^ F20 electron microscope(FEI, USA) operated at 80 kV to acquire morphological information of single phage particles.

### Thermal and acid-base stability tests

Thermal and acid-base stability tests were performed as previously described [18], with some modifications. The phage solution was diluted with SM buffer, after which the phage was treated at a specified temperature or pH for 1h. After treatment, the titer of the phage was determined by double-layer agar plate method. The results were expressed as phage stability in terms of the percentage of initial viral counts.

### Adsorption experiments and one-step growth analysis

Phage adsorption to host bacteria was performed as described previously [19], with minor modifications. The host strains were cultured to the logarithmic growth phase, mixed with an appropriate ratio of phage liquid. The mixture was incubated at 37 °C for and centrifuged immediately. The titer of the supernatant was determined using the method described above, and the phage adsorbed to the host bacteria was calculated based on the titer.

For the one-step growth analysis, the host bacteria was cultured to the logarithmic stage, and phage liquid was added at an appropriate MOI ratio of 10, incubated at 37 °C for 10 min, and centrifuged. The supernatant was discarded and the sediment was resuspended in liquid LB medium. The bacterial suspension was adjusted to 600 nm (OD_600_) of 0.12, and bacterial liquid was added to the LB liquid medium. The liquid was incubated with rotary shaking (200rpm, 37 °C), and aliquots of 100 µL were sampled from zero time to 1 h with 10 min intervals. The titer was determined using the double-layer agar plate method.

### Host range determination

The bacteria listed in Table 2 were used for host range analysis by standard spot tests [20]. Briefly, strains were grown overnight in LB medium. Suspension (10μL) purified phage suspension containing 10^7^ pfu/ml were spotted in the middle of a lawn of bacteria and left to dry before overnight incubation. Lysis characteristics were established at the spot where the phage was deposited. The assay host refers to the tested *K. pneumoniae* clinical isolates, and the isolation host refers to the *K. pneumoniae* isolate hvKpLS7, with which we initially isolated the phage.

### Sequencing and analysis of bacteriophage genomes

Bacteriophage DNA was prepared from high-titer phage preparations with phenol-chloroform and SDS [21]. DNA samples were sequenced in the second generation by Ion S5 (Thermo Fischer, USA) and third-generation by MinION (Oxford Nanopore Technologies, UK) genome sequencer. The sequencing data were assembled using the SPAdes assembler v. 3.13.2 [22], HGAP4 and Canu v1.6. MUMmer v3 [23] software was used to analyze the contigs obtained by splicing the second and third-generation sequencing data to reconfirm the assembly results and determe the positional relationship between the contigs, and to determine the gap between contigs.

To obtain information, the complete genomes of phages were sequenced and analyzed using a variety of bioinformatics tools. Open reading frames (ORFs) prediction were predicted using SoftBerry (http://www.softberry.com). Genome annotations were checked through sequence comparison of protein sequences using the blastn software (https://blast.ncbi.nlm.nih.gov). Genome comparative analysis was performed using Easyfig.

### Bacteriophage therapy assay and determination of virulence of strains

The *G. mellonella* model was used to evaluate the antibacterial efficacy of bacteriophages in *vivo* and the virulence of *K. pneumoniae* strains [24, 25]. The minimum lethal concentration of *K. pneumoniae* infection by larval caterpillars was determined to be 10^7^ CFU/mL within three days. When larvae did not respond to touch, they were considered dead. In the in *vivo* experiment, the larvae were divided into four groups, sixteen randomly chosen larvae were used for each group: (i) only injected with 10ul PBS, (ii) injected with 10ul of 10^6^ CFU/mL host bacteria, (iii) only injected with 10ul of 10^7^ PFU/mL of phage, (iv) injected with 10ul of 10^6^ CFU/ml host bacteria, and then injected with 10ul of 10^7^ PFU/mL phage within 30 min. All larvae were incubated at 37 °C and the number of dead larvae was counted at 12 h intervals up to 72 h after the incubation.

Virulence determination tests were performed according to the bacterial concentration in the previous phage treatment experiment (10^7^ CFU/mL). Sixteen *G. mellonella* larvae were injected with 10ul of the inoculum in every group. Survival was analyzed by Kaplan–Meier analysis with a log–rank test; differences were considered statistically significant at P < 0.05.

### Screen for phage-resistance strains

Phages hvKpP1 and hvKpP2 were mixed with the host bacteria for cultivation, and the mixture was cultured using double-layer soft agar. Plates were incubated overnight at 37 °C, and the resulting colonies were picked up and saved for further assays.

### Bacteria growth curves

All strains were cultured as described above. The following day, cultures were incubated in LB at a concentration of 1 × 10^7^ CFU/mL and added to individual wells of a 96-well microtiter plate. Plates were incubated for 12 h at 37°C and absorbance readings at 600 nm were recorded every 30 min using BMG SPECTROstar® Nano. Growth rates of the bacterial strains were calculated using three biological replicates.

### Mucoviscosity assay

The mucoviscosity of the capsule was assessed by low-speed centrifugation of the liquid culture [26]. Various overnight cultures of *K. pneumonia* were grown to adjust to OD_600_/mL of 1 and centrifuged at 1,000 ×g for 5 min. The OD_600_ values of the supernatants were then measured.

### Macrophage phagocytosis assay

RAW264.7 cells were seeded in 24-well plates and grown in Dulbecco’s modified Eagle’s medium (DMEM) (10% FBS, 100 mg/mL ampicillin and 100 mg/mL streptomycin) at 37 °C and 5% CO_2_. *K. pneumoniae* strains were added at a multiplicity of infection (MOI) of 10 bacteria per host cell and the inoculum was plated for colony forming units. Cells were rinsed three times with PBS and then incubated on fresh medium containing 300μg/ml gentamicin to kill extracellular bacteria. After three washes, cells were lysed with 0.1% Triton-X100 for 20 min, dilutions, and plated for bacterial CFU enumeration. The percentage of phagocytosed bacteria per inoculum was calculated and normalized to that of hvKpLS7. Three biological replicates per strain were used for each experiment.

### Transmission electron microscopy of bacteria

Transmission electron microscopy (TEM) was performed using JEM-1400PLUS (JEOL, Japan). Bacterial samples were perpared as described previously and modified [12]. Briefly, samples were fixed for at least 2 h at room temperature in 3% glutaraldehyde and post-fixed with 1% osmium tetroxide, dehydrated in alcohol grades, incubated with propylene oxide, and infiltrated overnight in a 1:1 mixture of propylene oxide and epoxy low-viscosity resin. The following day, samples were embedded on epoxy resin and polymerized. Ultrathin sections (approximately 50 nm) were cut on a Reichert EM UC7 microtome, transferred to copper grids stained with lead citrate, and examined using a JEM-1400PLUS transmission electron microscope, and images were recorded.

### Bacterial genome sequencing and analysis

Genomic DNA of wild-type hvKpLS7 and phage-resistant mutants was sequenced at Sangon Biotech (Shanghai, China) using the Illumina Hiseq platform (∼1 Gbp/sample, paired-end) as previously described [27]. The quality of raw sequencing reads was evaluated using FastQC. Low-quality reads and adapter sequences were trimmed using Trimmomatic software. Following the Genome Analyzer Toolkit (GATK) best practices pipeline, the genomic mapping tool Burrows-Wheeler Aligner (BWA) to map low-divergent sequences to the reference genome of *K. pneumoniae*. Mutations, including base substitutions, deletions, and insertions, were detected using SAMtools, MarkDuplicates, and BEDTools. DNA deletions mutations were further validated by PCR and sequencing.

### Cloning and complementation

Genomic DNA from hvKpLS7 was used as the template for wild-type gene cloning via PCR. PCR products were purified and cloned into the pBAD24-CM vector using the ClonExpress II One Step Cloning Kit (Vazyme, Nanjing, China). Recombinant plasmids were first heat-shocked into *E. coli* DH5α and further electroporated into corresponding phage-resistant mutants. The complementation strains were verified by PCR and sequenced using pBAD24 primers. The bacterial isolates transformed with an empty vector were tested in parallel.

### Statistical analysis

All experiments were performed with n equal to 3. Statistical analysis was performed using GraphPad Prism v.6.0 (Software Inc., La Jolla, CA, USA) to plot. For all macrophage phagocytosis and mucoviscosity assays, comparisons between mutant and WT, and WT and complementation strains were evaluated for statistical significance using an unpaired t-test. And the survival curves with the Kaplan-Meier method following a log-rank test to calculate the differences in survival. Statistical significance was set at P-value of < 0.05.

## Results

### Phage isolation and host range

Two lytic phages, vB_KpnP_cmc20191 (referred to as hvKpP1) and vB_KpnP_cmc20192 (hvKpP2), were isolated from sewage; microscopic observation of virion morphology by TEM showed that the phages were classified as members of the Podoviridae family (Fig 1A, B). They formed different plaques on the bacterial lawn of *K. pneumoniae* strain hvKpLS7 (Fig 1C, D). hvKpP1 produced clear plaques while hvKpP2 formed smaller lytic center plaques surrounded by a semi-transparent halo. The clinical host strain hvKpLS7 was characterized as an hvKP by 11 virulence-associated genes, including aerobactin, iroN, rmpA, rmpA2, ybtS, ureA,wabG, ycf, entB, iutA, and fimH [28] and was further confirmed in the *G. mellonella* model, using hypervirulent WCHKP030925 [29] as a positive control (Fig S1). This clinical strain belonging to K57 capsular serotype and host range experiments also confirmed that only strains belonging to K57 capsular serotype were specifically targeted by both phages, whereas others, including K1, K2, K64, K150, and K84 were not; it further confirmed that infections by phages had no relationship with MLST type (Table 1). K57 *Klebsiella pneumoniae* (K57-KP) was considered a highly virulent *Klebsiella pneumoniae* in clinical investigations [30, 31].

**Table 1.**
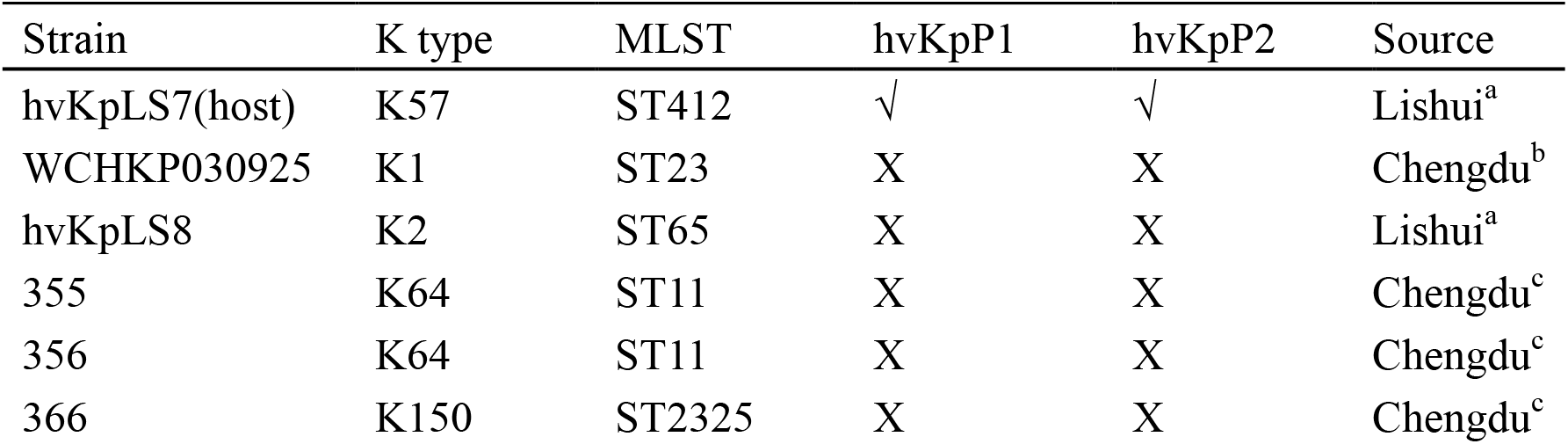
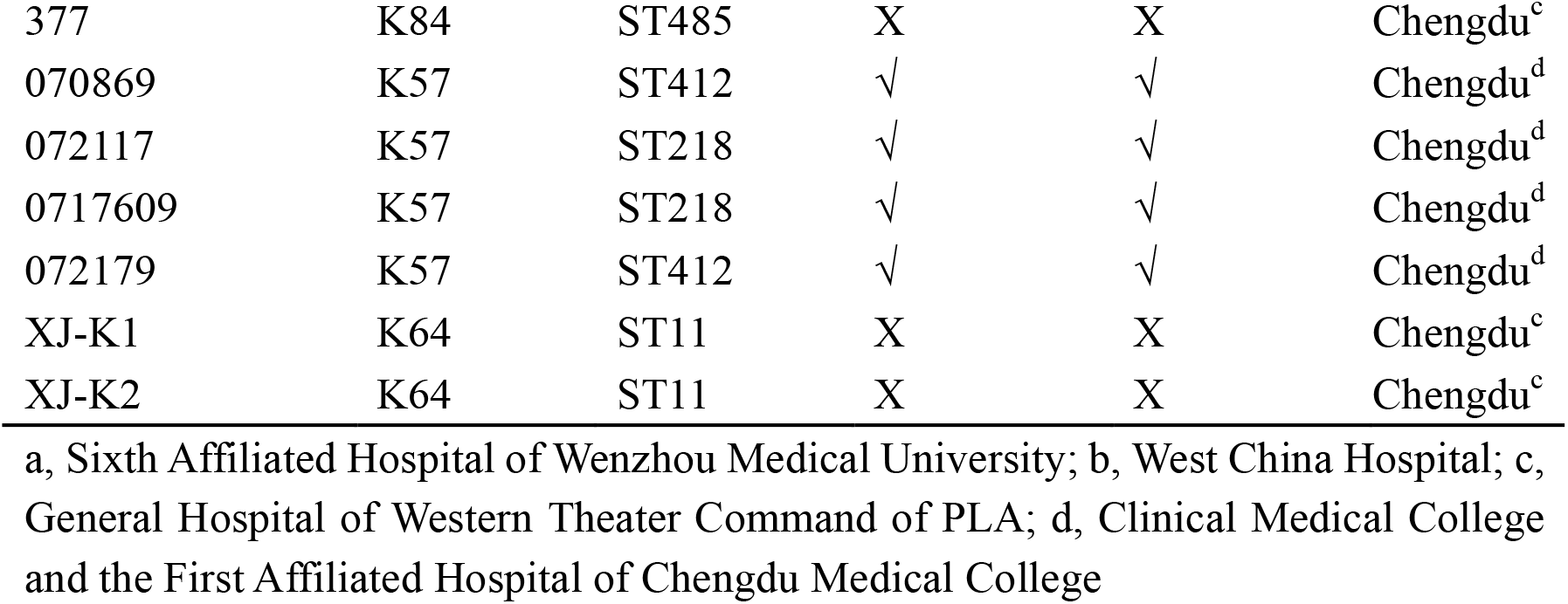
Host range of phages.

| Strain | K type | MLST | hvKpP1 | hvKpP2 | Source |
| --- | --- | --- | --- | --- | --- |
| hvKpLS7(host) | K57 | ST412 | √ | √ | Lishui<sup>a</sup> |
| WCHKP030925 | K1 | ST23 | X | X | Chengdu<sup>b</sup> |
| hvKpLS8 | K2 | ST65 | X | X | Lishui<sup>a</sup> |
| 355 | K64 | ST11 | X | X | Chengdu<sup>c</sup> |
| 356 | K64 | ST11 | X | X | Chengdu<sup>c</sup> |
| 366 | K150 | ST2325 | X | X | Chengdu<sup>c</sup> |
| 377 | K84 | ST485 | X | X | Chengdu<sup>c</sup> |
| 070869 | K57 | ST412 | √ | √ | Chengdu<sup>d</sup> |
| 072117 | K57 | ST218 | √ | √ | Chengdu<sup>d</sup> |
| 0717609 | K57 | ST218 | √ | √ | Chengdu<sup>d</sup> |
| 072179 | K57 | ST412 | √ | √ | Chengdu<sup>d</sup> |
| XJ-K1 | K64 | ST11 | X | X | Chengdu<sup>c</sup> |
| XJ-K2 | K64 | ST11 | X | X | Chengdu<sup>c</sup> |

a, Sixth Affiliated Hospital of Wenzhou Medical University; b, West China Hospital; c, General Hospital of Western Theater Command of PLA; d, Clinical Medical College and the First Affiliated Hospital of Chengdu Medical College

**Fig 1.**
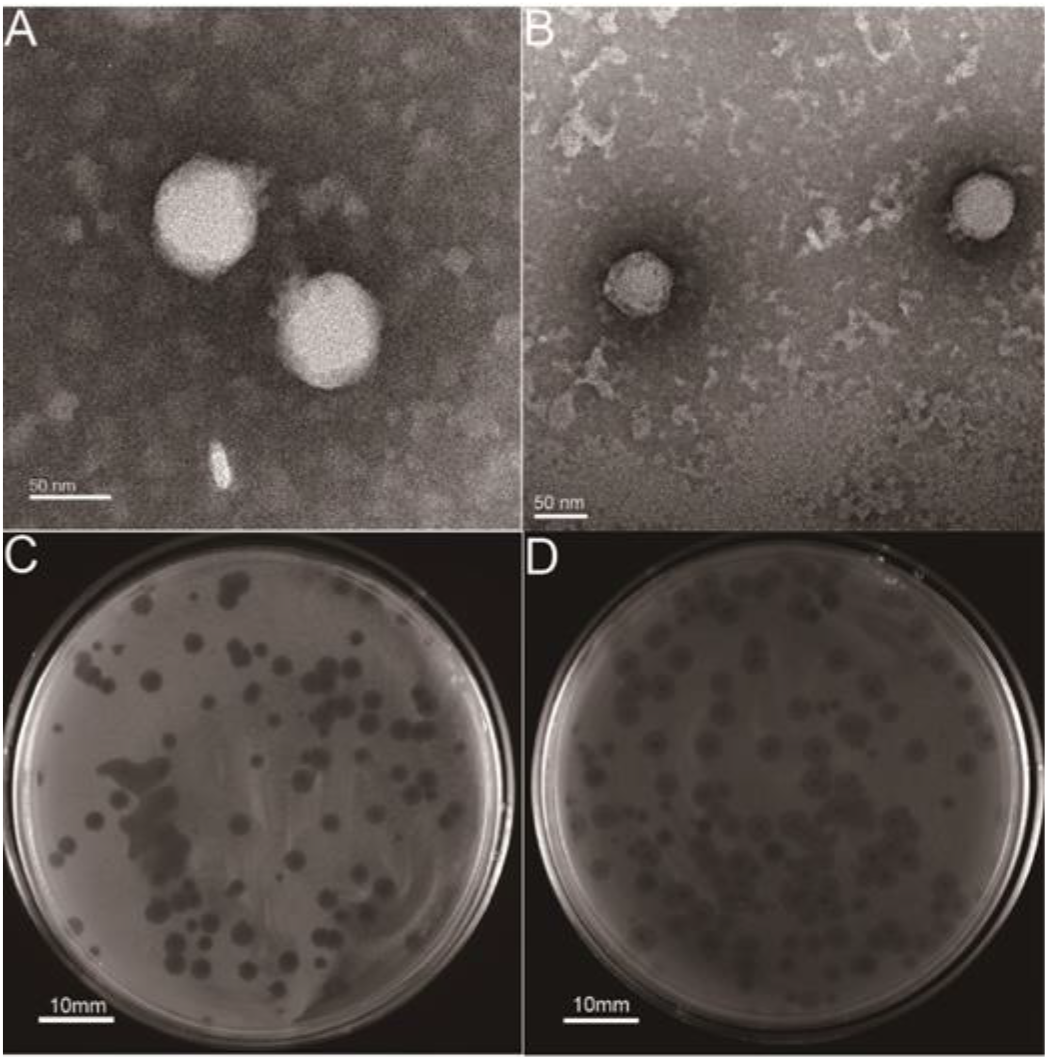
Structural characterization and plaque morphology of isolated phages. (A), (B) Transmission electron micrograph (TEM) of phage hvKpP1 and hvKpP2, Bar = 50 nm. (C), (D) Plaques of phage hvKpP1 and hvKpP2 on *K. pneumoniae* hvKpLS7, Bar = 10 mm.

### Phage characterization

To determine phage stability, the sensitivity of phages to temperature and pH stability was analyzed (Table 2). Phages were stable in the pH range of 4-11 or under 70°C (Fig S2AB). These results were in line with those of previous studies [32, 33], showing that these two phages could maintain high lytic activity under broad physicochemical conditions. The adsorption rate curves of the two phages showed that more than 90% of bacteriophages were adsorbed within 10 min (Fig S2C). According to the one-step growth curves (Fig S2D), the replication cycle of hvKpP2 was approximately 60 min. However, the latent period of hvKpP1 was relatively short, only approximately 30 min. The burst size of hvKpP1 (149 PFU/Cell) was greater than that of hvKpP2 (96 PFU/Cell). The eclipse period and burst size may be part of the reason for the difference in plaque production between the two phages. A summary comparison of these two phages is presented in Table 2.

**Table 2.**
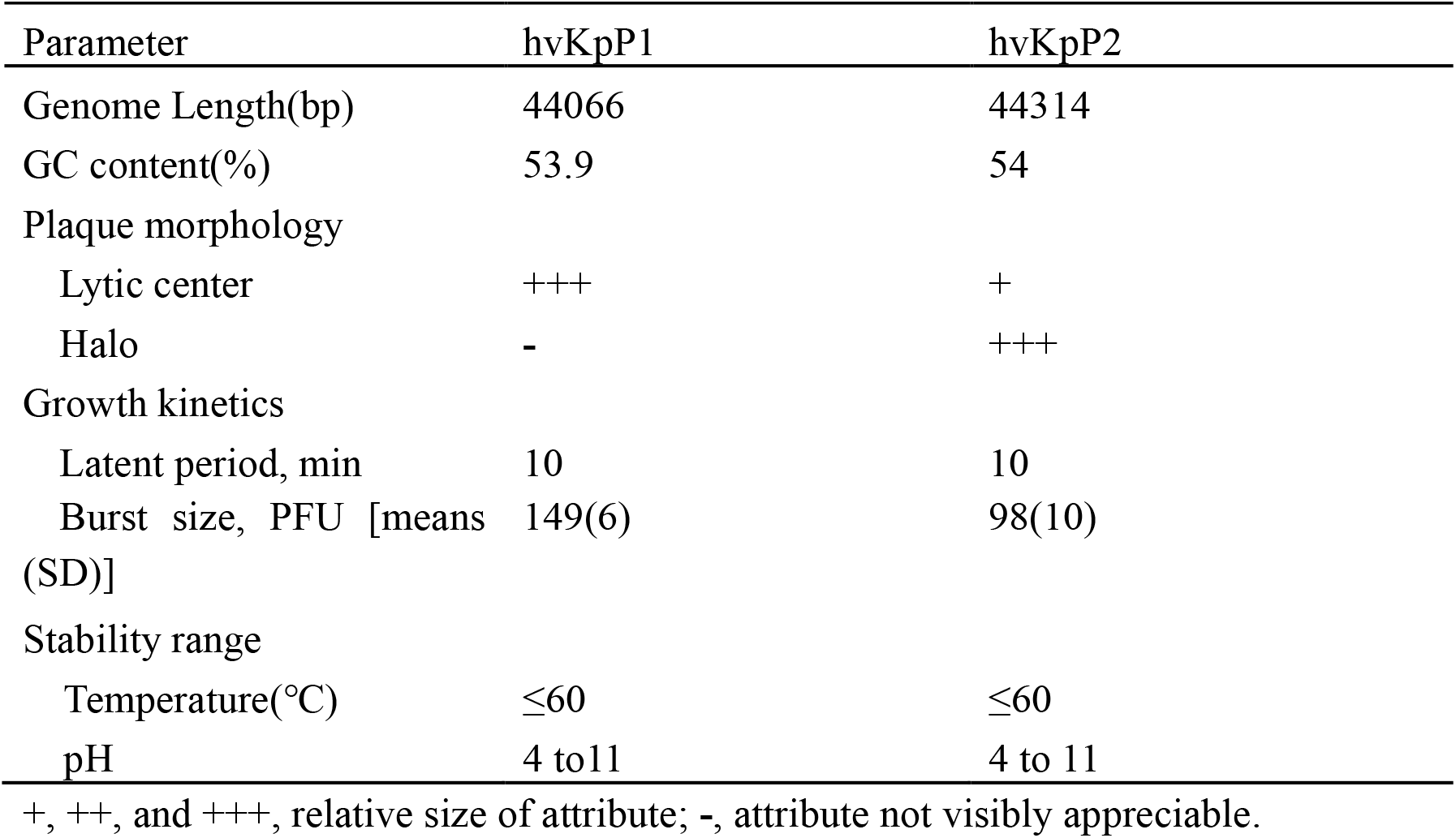
Summary phage comparison.

| Parameter | hvKpP1 | hvKpP2 |
| --- | --- | --- |
| Genome Length(bp) | 44066 | 44314 |
| GC content(%) | 53.9 | 54 |
| Plaque morphology |  |  |
| Lytic center | +++ | + |
| Halo | - | +++ |
| Growth kinetics |  |  |
| Latent period, min | 10 | 10 |
| Burst size, PFU [means (SD)] | 149(6) | 98(10) |
| Stability range |  |  |
| Temperature(°C) | ≤60 | ≤60 |
| pH | 4 to 11 | 4 to 11 |

+, ++, and +++, relative size of attribute; -, attribute not visibly appreciable.

Bioinformatics analysis helps us to better understand and predict biological characteristics of phages. Thus, the complete genomes of the two phages were sequenced and analyzed using bioinformatics tools. The genome of phage hvKpP1 was 44066 bp in length with 54% G+C content, while hvKpP2 was 44314 bp in length with 53.9% G+C content. In total, 55 and 58 putative coding regions (CDSs) were detected by RAST analysis. Importantly, the lack of lysogeny, host conversion, and toxins supported the growth kinetics data, suggesting that the two phages possess lytic properties and could be used for therapeutic purposes [34]. Nucleotide BLAST analysis revealed that hvKpP1 and hvKpP2 exhibited high DNA similarity. We compared their genomes using Easyfig. As shown in Fig 2, the main difference between hvKpP1 and hvKpP2 lies in the genes encoding the DNA packaging. Among them, hvKpP1 contains more genes encoding HNH family proteins, which are considered to be related to phage DNA replication in reference research [35]. This may be the main reason why hvKpP1 bursted more than hvKpP2. The two phages had high homology in the tail packaging region, indicating that their adsorption targets for the host bacteria were the the same. The complete nucleotide sequences of phages vB_KpnP_cmc20191 and vB_KpnP_cmc20192 were determined and deposited in GenBank under the accession numbers MT559526 and MT559527, respectively.

**Fig 2.**
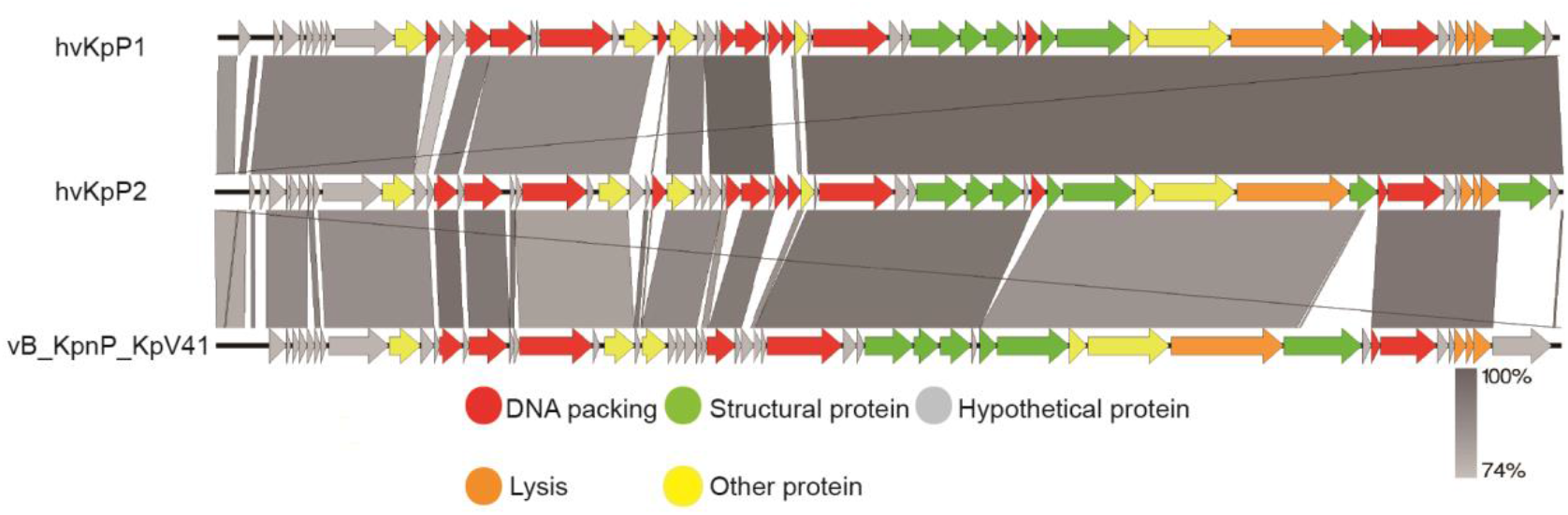
Pairwise BLASTn comparison and phylogenetic tree of hvKpP1, hvKpP2. The genome map was performed using the Easyfig. Arrows represent predicted ORFs, the direction of the arrow represents the direction of transcription. Different colors denote different functional groups of bacteriophage genes.

### In *vivo* efficiency of bacteriophage treatment

The efficacy of phages hvKpP1 and hvKpP2 was evaluated in *vivo* using the *G*.*mellonella* larvae model. For larva infected by the host strain hvKpLS7, the survival rate was only 5 and 10 % in three days, respectively. In the phage treatment groups, survival was significantly superior, with three-day survival rates of 75 and 90%. There was a significant difference in the survival rates between the larva infection group and the treatment group (P<0.05). Additionally, the larvae group injected only with phages still had a high survival rate, demonstrating the safety of the phages in this model (Fig 3). The present study reports, for the first time, on phage efficacy against K57 capsular serotype *K. pneumoniae* in *G. mellonella*.

**Fig 3.**
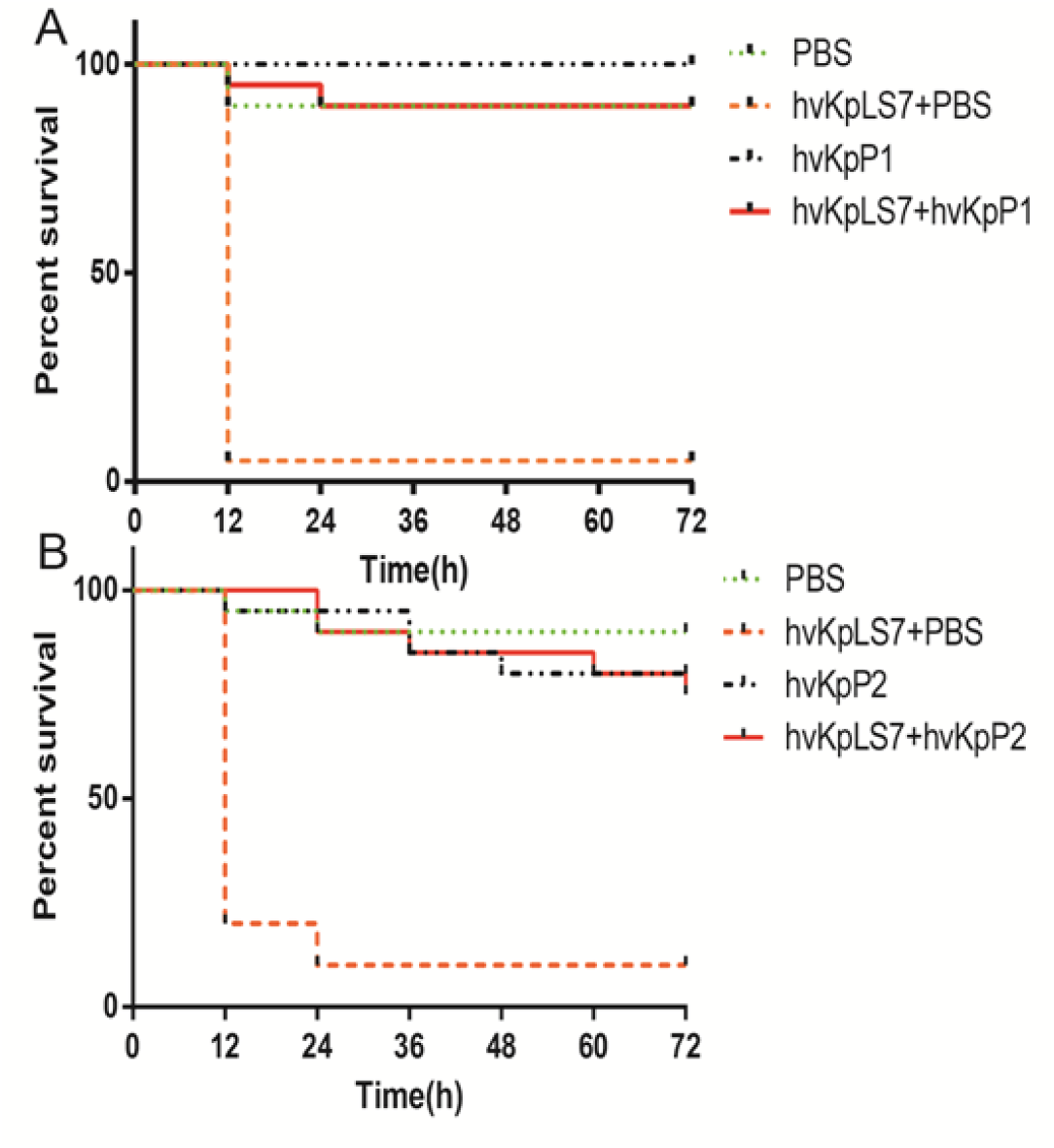
Survival of *G. mellonella* larvae after infection by hypervirulent *K. pneumoniae* hvKpLS7 and phages therapy. (A) Treatment with hvKpP1 (B) Treatment with hvKpP2.

### Virulence declined with phage-resistance *K. pneumonia*

Although phage treatment in the *G. mellonella* model showed excellent results, phage-resistant *K. pneumoniae* colonies were easily produced on bacterium-phage-co-cultured LB agar plates. Interestingly, phage-resistant *K. pneumonia* did not affect therapy. To test the difference between wild type and phage-resistant *K. pneumoniae*, six mutants were randomly selected from the plates. We then evaluated whether phage-resistant bacteria changed their generation time, and the results indicated that resistance to phage had no effect on growth (Fig 4B). On the LB agar plates, the wild-type host bacteriaium hvKpLS7 was moist, hypermucoid and was reflective when photographed, whereas the phage-resistant bacteria had a translucent appearance and had a reduced ability to produce mucoid (Fig 4A). And all the mutant strains shown negative string test results (results are not shown). These changes were similar to those reported by Tan et al. [27]. Some studies have suggested that HMV is related to the capsule [36], hence, mucoviscosity of the capsule was assessed by low-speed centrifugation of liquid culture. Strains were grown in LB, diluted to an optical density at 600 nm (OD_600_) of 1, and then subjected to low-speed centrifugation. During the centrifugation process, hvKpLS7 did not sediment well, the supernatant was still turbid, and the OD_600_ of the supernatant was 0.35. Contrastingly, the phage-resistant bacteria was well precipitated in the low-speed centrifugation experiment, the supernatant was relatively clear, and the average OD_600_ decreased to 1/3, compared to the wild type (Fig 4CD). This indicated that *K. pneumoniae* with phage resistance significantly decreased capsular adhesion.

**Fig 4.**
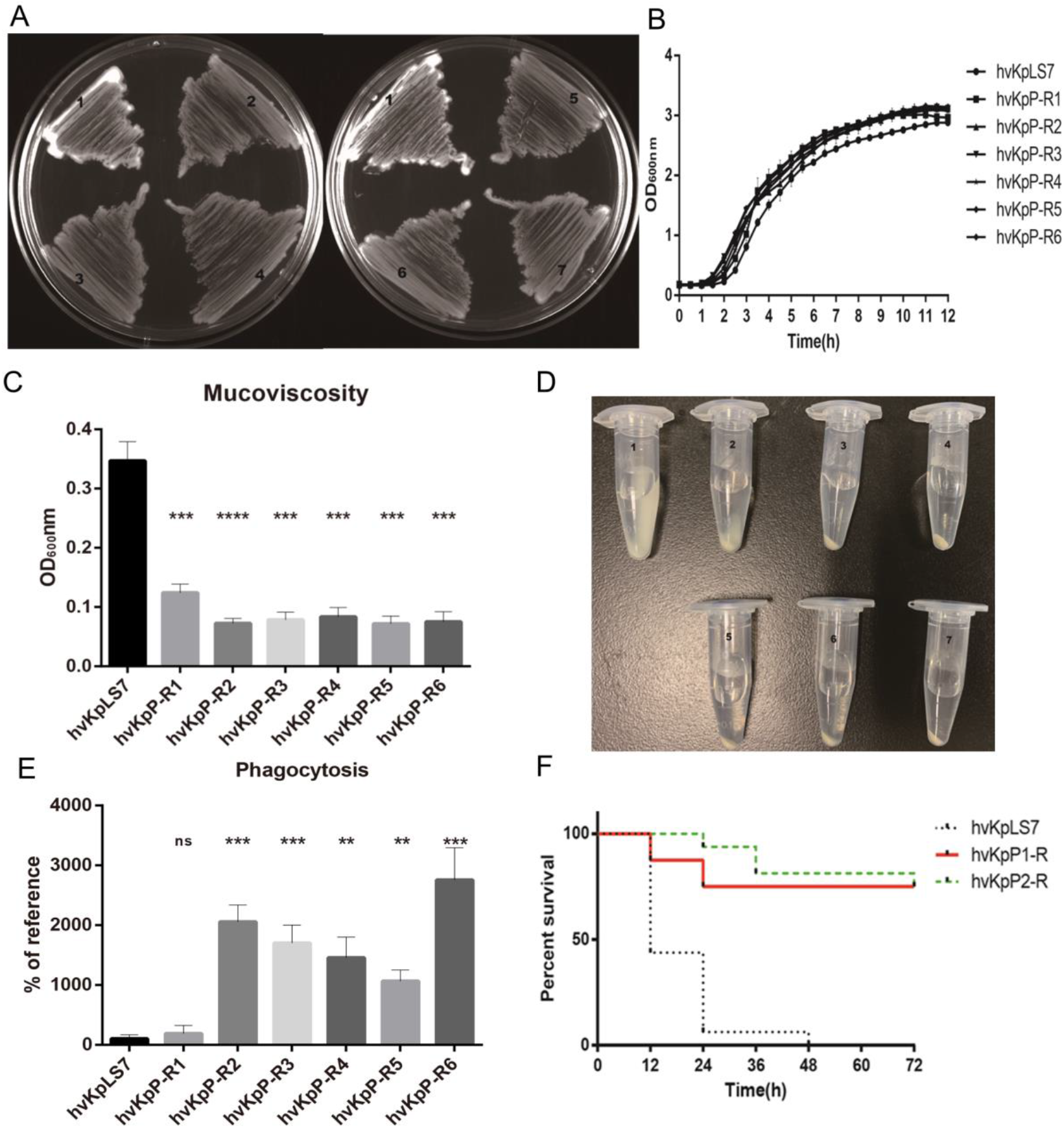
The virulence changed with *K. pneumoniae* hvKpLS7 and hvKpP-R. (A) Morphological comparison between colonies of *K. pneumoniae* on LB-ager plate. hvKpLS7 (mucoid, moist, and sticky), and phage-resistant mutant (dry, rough, and transparent). (B) Growth curves of the wild-type and mutant of *K. pneumoniae* strains. (C), (D) Centrifugation analysis of hvKpLS7 and hvKpP-R. (E) Phagocytosis by RAW264.7 macrophages. (F) Survival of *G. mellonella* larvae. The unpaired t-test was performed to determine statistically significant differences between each mutant and hvKpLS7. **, P ≤ 0.01 ***, P ≤ 0.001; ****, P ≤ 0.0001; ns, not significant.

Previous studies have demonstrated that *K. pneumoniae* capsules confer significant phagocytosis resistance to macrophages [37]. These strains were co-incubated with phagocytes RAW264.7, the lysates of washed phagocytes were daubed on agar plates, and the bacterial colonies were counted and recorded. Results showed that amount of the mutant strain (except hvKpP-R1) devoured by the phagocytes increased at least 10 times compared to that of the wild-type, indicating that the phagocytes could effectively eliminate these mutant bacteria.

Further, whether the resistance of hvkp to phage affects its virulence during infection was determined in *vivo*. As expected, data of the infection with *G. mellonella* showed that the phage-resistant *K. pneumoniae* had a higher survival rate with the same concentration of bacteria (Fig 4F). The results of these assays indicate that the phage-resistant *K. pneumoniae* reduced their virulence, which might be due to the absence of a capsule.

### Identification of mutant genes in phage resistant strains

To identify the genes responsible for resistance in bacteria, genomes of wild-type hvKpLS7 and phage-resistant mutants were sequenced using the Illumina Hiseq platform and comparatively analyzed. High-probability mutations (defined as high-frequency, non-silent mutations within an open reading frame) were selected for further validation. These mutant strains had, mutations in two genes associated with the capsule, *wzc* and *wcaJ*, which corresponded to hvKpP-R2 and hvKpP-R3 respectively. Previous studies have shown that hvkp contains a compact gene cluster, which plays an important role in capsule synthesis.

### Complementation of strain hvKpP-R2 with wild-type *wzc* restore phage sensitivity and virulence

The *Wzc* gene belongs to the capsular polysaccharide synthesis gene clusters, which facilitates polymerization of capsular polysaccharide when activated by its tyrosine kinase(TK) domain [38]. As shown in Fig 5A, only one base was deleted in the *wzc* gene of the mutant strain hvKpP-R2. The deletion of 1463 base A resulted in a frameshift of the *wzc* gene, therefore mutation resulted in missing TK domain and loss of catalytic activity. To confirm that the mutation complementation of strain hvKpP-R2 with wild-type *wzc* restored the sensitivity and mucoid phenotypes of the mutant strains, we amplified and cloned the *K. pneumoniae* strain hvkpLS7 *wzc* into plasmid vector pBad24.

**Fig 5.**
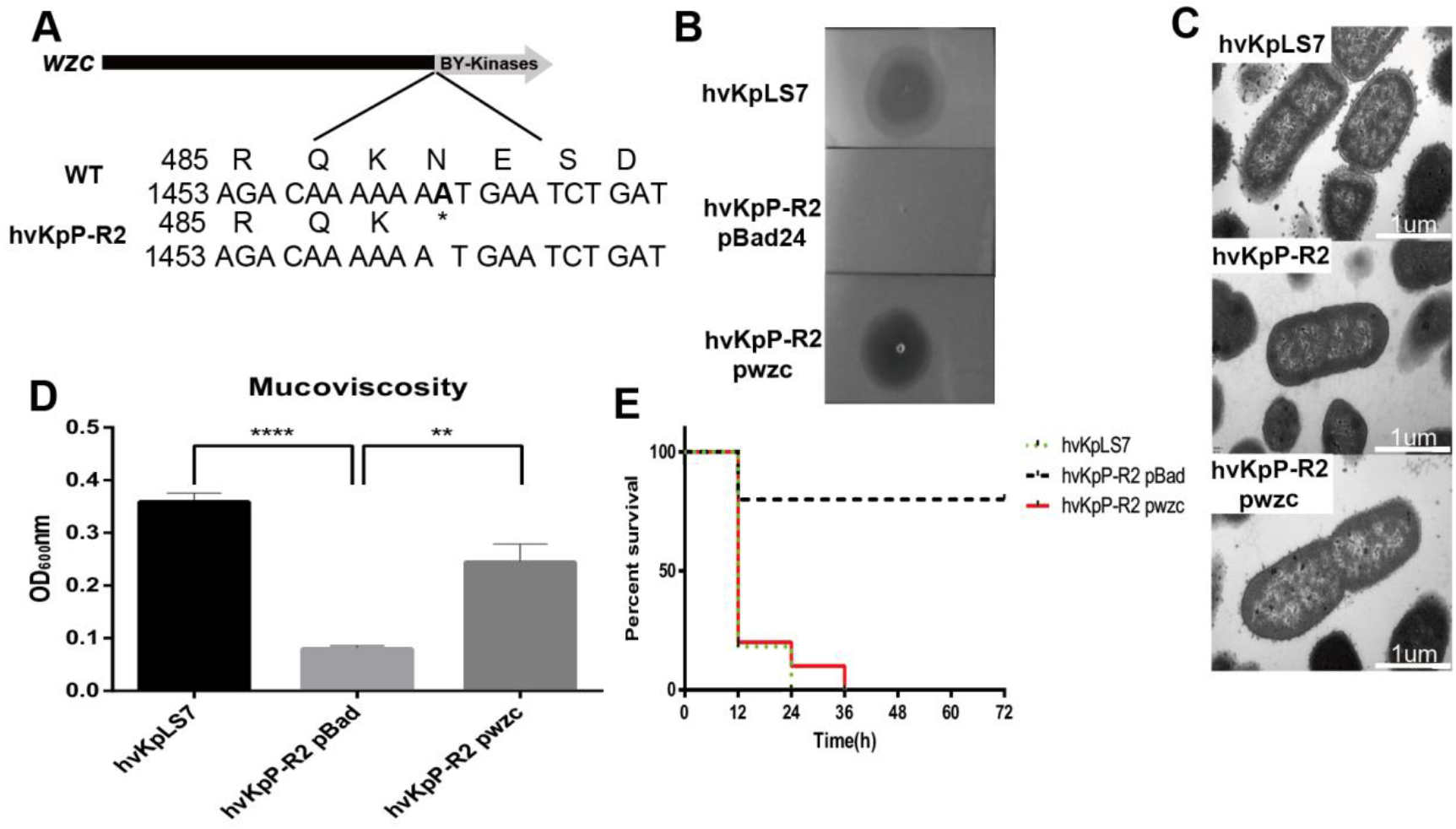
Complementation of strain hvKpP-R with wild-type *wzc* restore phage sensitivity and virulence. (A) Schematic representation of *wzc* in *K. pneumoniae* hvKpLS7 and hvKpP-R2. Genes are represented as arrows. (B) Spot test assay of phage on the parental *K. pneumoniae* strain hvKpLS7 and its derived mutants (hvKpP-R2 pBad and hvKpP-R2 pwzc). (C) The *wzc* gene affects capsule production. TEM of wild-type, phage-resistant and complementation strain. For every isolate, one representative image from six images obtained from one section is shown. (D) Mucoviscosity is restored in hvKpP-R2 pwzc (**, P ≤ 0.01; ****, P ≤ 0.0001). (E) Survival rates. Survival rates of *G. mellonella* with hvKpLS7, hvKpP-R2 pBad and hvKpP-R pwzc were determined.

Spot test results indicated that the strain hvKpP-R2 pwzc restored phage sensitivity (Fig 5B). Low-speed centrifugation of the liquid culture showed increased mucoviscosity of the capsule (Fig 5D). TEM analysis showed that the boundaries of hvKpP-R2 were smoother than those of LS7 and hvKpP-R2pwzc, which confirmed that phage resistance selection caused the loss of the hvkp capsule, and the expression of recombinant *wzc* could significantly restore the polymerization of capsule(Fig 5C). According to the three-day survival rate of *G. mellonella*, results of the hvKpP-R2pwzc and wild-type infection groups were similar (Fig 5E), indicative that after complementing the *wzc* gene of the wild-type strain, the virulence of this mutant strain was rescued.

Therefore, after a mutation in *wzc*, the ability of *K. pneumoniae* to synthesize the capsule was limited, and phages could not adsorb the mutant strain. Similarly, the virulence of the mutant strain decreased due to the loss of the capsule. Previous studies have shown that since *wzc* plays an indispensable role in capsular polysaccharide surface assembly in *K. pneumoniae*, it is conceivable that mutations in *wzc* are dominant in phage-resistant strains [39].

### Complementation of strain hvKpP-R3 with wild-Type *wcaJ* restore phage sensitivity virulence

We amplified and cloned *wcaJ* from wild-type to phage-resistant mutant hvKpP-R3, whose *wcaJ*, compared with the wild-type, had four bases deleted (Fig 6A). As described above, the same experiments were carried out on hvKpP-R3 pwcaJ, and results are shown in Fig 6B. The phage-resistant hvKpP-R3 containing the wild-type *wcaJ* gene can be re-infected by the two phages, and the adhesion of the bacteria was restored to a certain extent. Electron microscopy images (Fig 6C) showed no differences in cell size and shape between the wild-type and phage-resistant mutants. The most evident difference was the loss of capsular in the mutant strain. Replenishment of the *wcaJ* gene allowed the mutant strain to regenerate the capsule, similar to the *wbap* gene [12]. In the in *vivo* infection experiment of *G. mellonella*, although the lethality rate of the hvKpP-R3 pwcaJ was lower than that of the wild-type in the first 12 h, it was significantly higher than that of the mutant strain hvKpP-R3, and within three days was also comparable to that of the wild type (Fig 6E).

**Fig 6.**
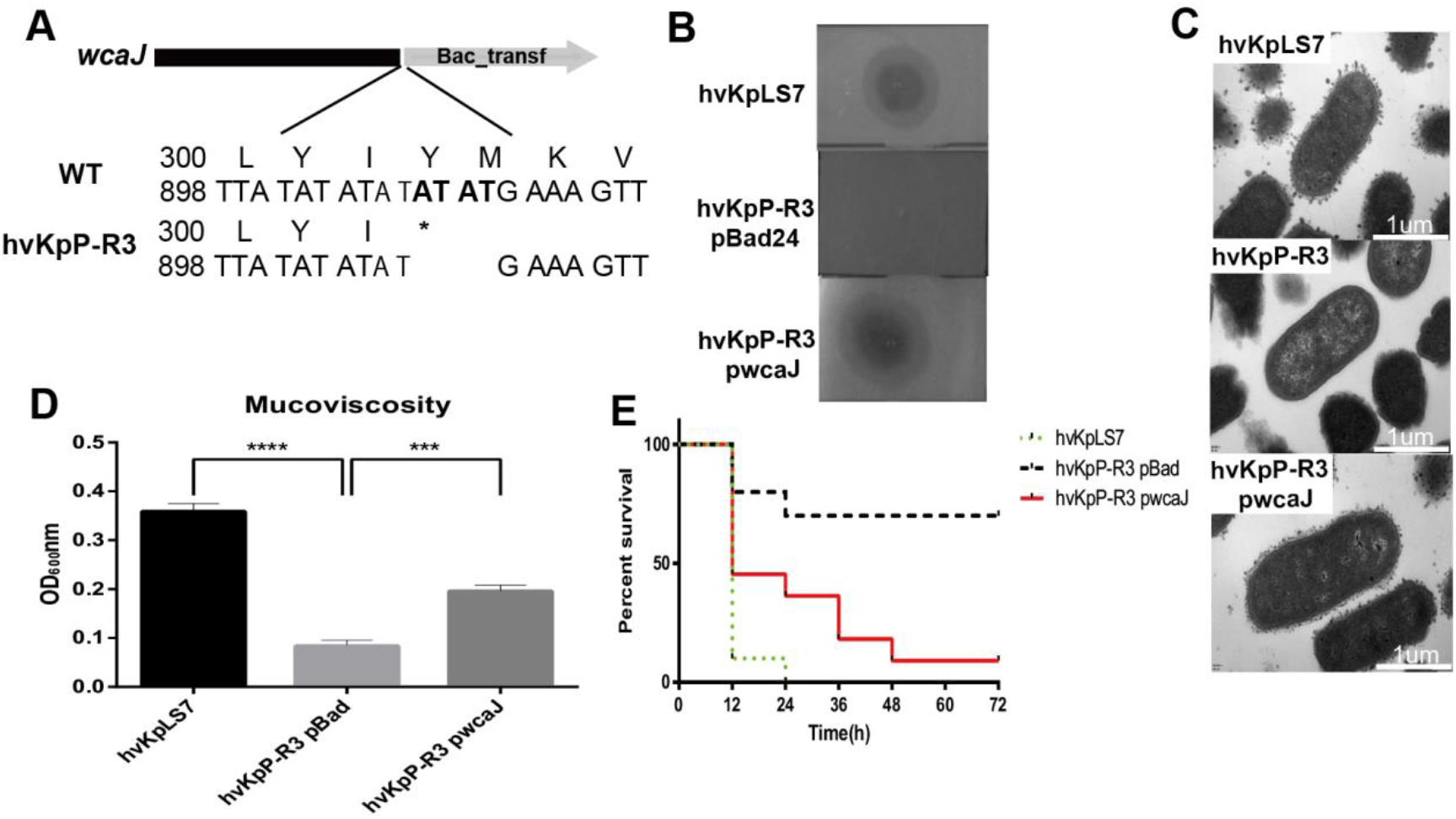
Complementation of strain hvKpP-R with wild-type *wcaJ* restore phage sensitivity and virulence. (A) WT, wild-type;Schematic representation of *wcaJ* in wild-type *K. pneumoniae* and hvKpP-R3. Genes are represented as arrows. (B) Spot test assay of phage on the parental *K. pneumoniae* strain hvKpLS7 and its derived mutants (hvKpP-R3 pBad and hvKpP-R3 pwcaJ). (C) The *wcaJ* gene affects capsule production. TEM of wild-type, phage-resistant and complementation strain. For every isolate, one representative image from six images obtained from one section is shown. (D) Mucoviscosity is restored in hvKpP-R3 pwcaJ (***, P ≤ 0.001; ****, P ≤ 0.0001). (E) Survival rates. Survival rates of *G. mellonella* with hvKpLS7, hvKpP-R3 pBad and hvKpP-R pwcaJ were determined.

As a gene present in *K. pneumoniae cps*(capsular polysaccharide*), wcaJ* was the first glycosyltransferase (GT) to transfer sugar to the lipid acceptor and initiates the synthesis of the capsular repeat [40], therefore, strains of *wcaJ* mutation could not produce the capsule. Although the phenotypes of the mutant strain were similar to those of the Tan et al [27] study, it was different from the mutant strain in that a 770 bp insertion sequence derived was detected between the *wcaJ* gene, while the mutant hvKpP-R3 in this study only had four base deletions in *wcaJ*, indicating that the mutation generation approach is diverse.

## Discussion

Hypervirulent *K. pneumoniae* usually infects healthy individuals in the community. In recent years, the emergence of carbapenem-resistant hypervirulent *K. pneumoniae* has posed regarded as a serious threat to public health. In this era of emerging antibiotic resistance, phage therapy is resurfacing as a conceivable alternative for treatment, which is considered a cost-effective alternative to antibiotics. Although phages have been used clinically [41, 42], a detailed understanding of phage biology is required. In this study, two podoviridae phages, hvKpP1 and hvKpP2, were isolated and characterized. The phages identified in the present study differed from other *K. pneumonia*e phages found in previous studies [32, 43, 44] in that they could lyse the K57 capsular, type hypervirulent *K. pneumonia*e, within a short lytic life cycle, producing a large burst size. Bioinformatics analysis showed that these phages do not contain lysogeny, host conversion, or toxin genes, proving that the newly discovered phages can be used for treatment.

During phage therapy, the host can develop resistance to the phage using various mechanisms [45]. Phage-resistant bacteria have been previously reported [46, 47], and in this study, phage-resistant mutations were easily manipulated in vitro. Results showed that the phage therapeutic effect was not affected by these mutants, and instead the virulence of these mutants was significantly reduced. This prompted a further investigation on the mechanisms of phage resistance and virulence reduction. Mutants were randomly selected on bacteria-phage co-cultured plates the colonies produced by these mutants were smaller and rougher than those of the wild type bacteria. Based on previous studies [48] these changes were attributed to the decreased ability of the bacteria to produce capsules. Bacterial capsular polysaccharides are produced by different *Enterobacteriaceae* strains, which play pivotal roles in bacterial survival under different environments and during host infection [49]. As one of the factors contributing to the hypervirulent *K. pneumoniae*, the loss of the capsule leads to changes in virulence [50, 51]. In this study, the virulence of the phage-resistant mutant strain was significantly reduced; this was at the cost of phage adsorption as the bacteria could not be absorbed by the phage after losing the capsule.

Genome sequencing of the screened bacteriophage-resistant bacteria revealed a base deletion in two genes, *wzc* and *wcaJ*, related to capsule production. Both genes form part of the capsule synthesis gene cluster [52], which encode tyrosine-phosphorylated proteins, and are speculated to be involved in the converging signal. From the results of this study it was evident that both gene mutations affected the production of the capsule, leading to an interruption of phage adsorption. The *wzc* of *E*.*coli* phosphorylates an endogenous UDP-glucose dehydrogenase (Ugd) involved in the production of exopolysaccharide colanic acid, as well as the production of UDP-4-amino-4-deoxy-L-arabinose. Capsular polysaccharides are synthesized by the *wza-wzb-wzc* system, which enhances the virulence of bacteria [53]. Hesse et al. [39] selected 57 strains of phage-resistant mutants from plates and broth cultures, and the mutated parts of these bacteria were mainly related to capsular synthesis and LPS, among which *wzc* produced the most mutated resistant bacteria. In the present study, the *wzc* mutation significantly reduced the virulence of *K. pneumoniae*.

Lin et al. made site-specific mutants in *wcaJ*, replacing each indicated amino acid (tyrosine) with phenylalanine coding sequences. *K. pneumoniae*, which had genetic mutations, exhibited a reduction in colony size, decreased mucosity, and reduced virulence [54]. Cai et al. [50]deleted the entire single gene *wcaJ* of capsular polysaccharide, which phages could not infect, and the colony morphology was small and rough, resembling phage-resistant bacteria. Mutation found in this study achieved the same effect by only a few base deletions, strengthening our understanding of the underlying mechanisms to acquired resistance in bacteria. Our data showed that complementing *wzc* and *wcaJ* genes into the mutant strains enabled the strains to re-infect and regain virulence.

This study only detected that hypervirulent *K. pneumoniae* with mutations in two genes related to capsule production could develop resistance to the new phages while having reduced their virulence. Whether mutations in other genes have the same effect requires further research.

## Conclusions

Taken together, the two new bacteriophages isolated have the ability to kill K57 capsular hypervirulent *K. pneumoniae*. In the *in vivo* experiment, the bacteriophages were effective in treating bacterial infections in the *G. mellonella* larvae model. This study also confirmed that the virulence of phage-resistant bacteria decreased as they developed resistance to phages. Through gene sequencing analysis and complement experiments, it was further verified that one or several base deletions of *wzc* and *wcaJ* genes played a role in phage receptor loss, resulting in no adsorption by the two phage strains, and reducing virulence at the same time. Our results suggest that hvKpP1 and hvKpP2 can be promising candidates for further investigation in phage-therapy research. As these phages targeted a hypervirulent serotype, and all their examined properties were suitable, our results may aid the development of bacteriophage-based therapeutic strategies for *Klebsiella pneumoniae* infections, specifically targeting hypervirulent strains.

## Acknowledgments

We are grateful to Dr. Zhiyong Zong of West China Hospital for the gift of WCHKP030925.

## Supporting information

**S1 Fig.**
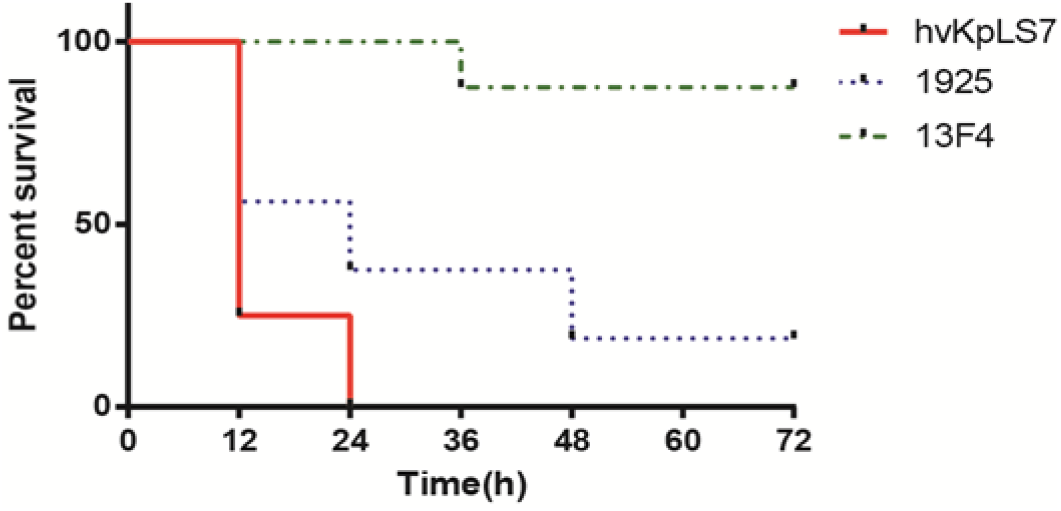
Survival of *G. mellonella* after infection with *K. pneumoniae* strains. The effect of 1 × 10^7^ CFU/mL of each strain on survival of *G. mellonella* is shown. hvKpLS7 was the host of phages, while 1925 and 13F4 were used as controls.

**S2 Fig.**
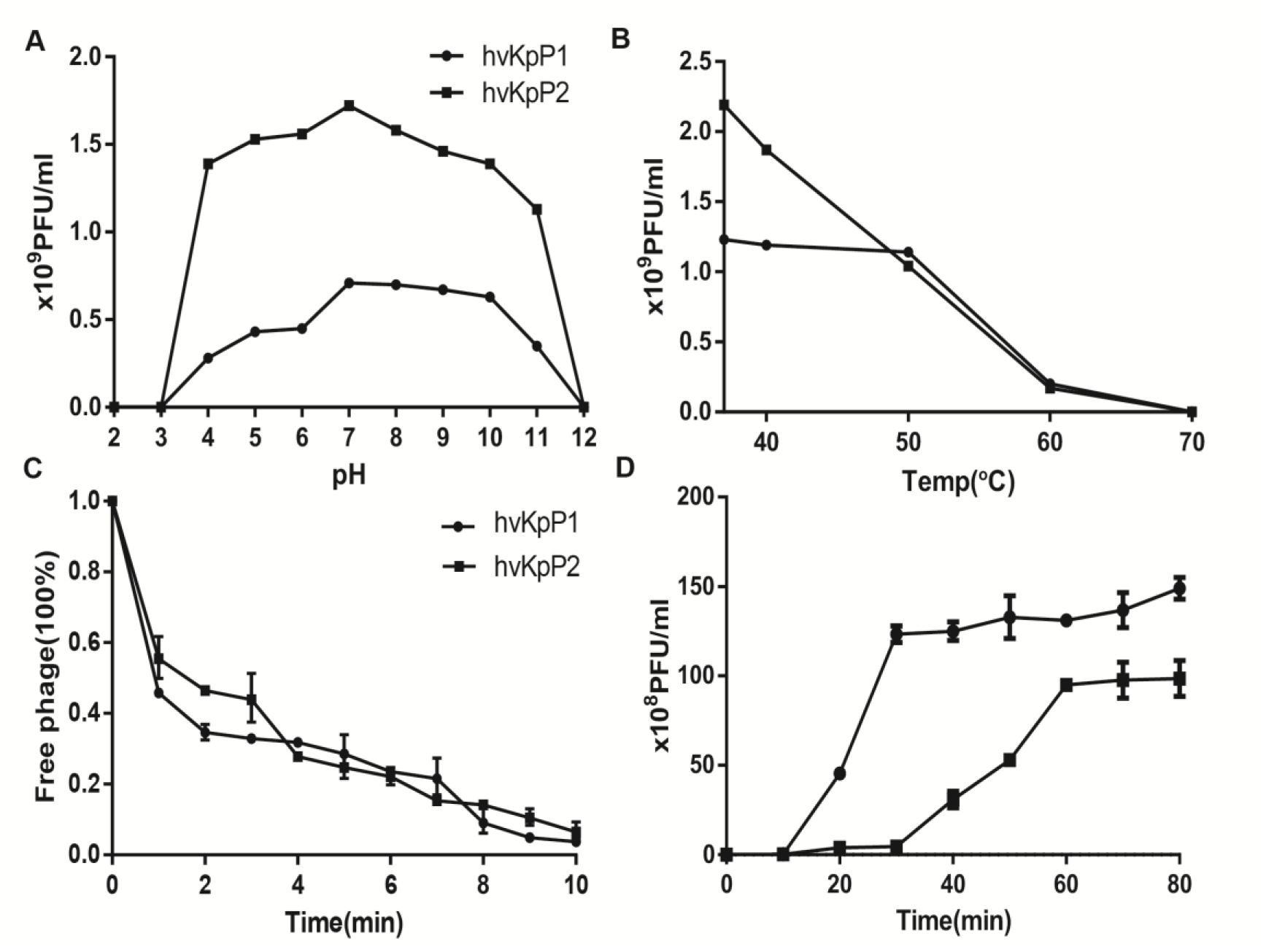
Biological characterization of phages. A, Survival rate of phages at different temperatures. B, Phage stability at variable pH. C, Adsorption rate. D, One-step growth curve.

